# Two-step polar plastid migration via F-actin and microtubules ensures unequal inheritance during asymmetric division of Arabidopsis zygote

**DOI:** 10.64898/2026.01.19.697633

**Authors:** Keigo Tada, Hikari Matsumoto, Takao Oi, Zichen Kang, Tomonobu Nonoyama, Satoru Tsugawa, Yusuke Kimata, Shuhei Kusano, Shinya Hagihara, Shintaro Ichikawa, Yutaka Kodama, Minako Ueda

## Abstract

The zygote is the origin of development, and in most angiosperms, it divides asymmetrically to establish the apical–basal axis. In *Arabidopsis thaliana*, various organelles in the zygote undergo polar migration along actin filaments (F-actin), resulting in unequal inheritance, but the behavior of plastids, essential precursors of chloroplasts, has remained unclear. Here, using quantitative live-cell imaging, we reveal that plastids undergo two-step polar migration: they first move apically together with the nucleus along F-actin, and when nuclear migration slows, they switch to microtubule (MT)-dependent migration to move further apically. This results in unequal plastid inheritance by the apical cell. Although these plastids are amyloplasts containing starch granules, starch is dispensable for migration, unlike the gravity response. Instead, a fertilization-activated MAP kinase pathway is required for polar plastid migration. Our results demonstrate that the zygote possesses a spatiotemporal regulatory mechanism that ensures unequal plastid inheritance at the onset of plant ontogeny.

## Introduction

In various plant species, the multicellular plant body arises from a single zygote. In *Arabidopsis thaliana* (Arabidopsis), the first asymmetric division of the zygote establishes the apical–basal axis, with a small apical daughter cell that extensively proliferates to form most of the plant body and a larger basal daughter cell that gives rise to the extra-embryonic suspensor ^1,2^. Live-cell imaging of zygotes has revealed that mitochondria and vacuoles migrate along longitudinal F-actin cables and become densely distributed toward the apical and basal cells, respectively ^3-5^. Disruption of this unequal inheritance causes defective embryo patterning; for example, vacuoles that are aberrantly inherited by the apical cell persist in its cell lineage and impair organogenesis, indicating that precise control of organelle positioning at the time of zygotic division is critical for proper embryogenesis ^5^.

Plastids contain their own genome and cannot be synthesized *de novo*, so they must be properly segregated at each cell division beginning with the first zygotic division to be transmitted through all cell lineages. Egg cells and zygotes in various plant species, including cotton and rice, harbor large numbers of plastids ^6,7^, and Arabidopsis mutants with impaired plastid function arrest development at the first or second zygotic division ^8,9^. Despite these shared features and their essential role, plastid dynamics in living zygotes and their relationship to axis formation have remained unclear. We therefore set out to characterize the spatiotemporal behavior of plastids in Arabidopsis zygotes by combining quantitative live-cell imaging, electron microscopy, fluorescent probes and pharmacological and genetic perturbations.

## Results

### Plastids migrate in two steps toward the apical tip of the zygote

We analyzed the plastid dynamics in Arabidopsis zygotes using time-lapse observations with a plastid/nucleus/PM marker and two-photon excitation microscopy (2PEM) (Fig. 1a, b and Video S1). This marker includes a plastid marker (EC1p::TP-Clover), a nucleus marker (ABI4p::H2B-tdTomato), and a plasma membrane (PM) marker (EC1p::RRvT-SYP121) (see Methods). In unfertilized egg cells, plastids extended filamentous structures, i.e., stromules, and occupied a position just below the apically located nucleus (Fig. 1a, c), consistent with electron-microscopy observations of non-transgenic plants ^10^. After fertilization (00:00 in Fig. 1b, c), cell shrinkage occurred as previously reported ^3^, and plastids surrounded the nucleus at the cell center. During subsequent zygote elongation, plastids migrated apically together with the nucleus, then adopted an elongated shape and passed along the side of the nucleus to move beyond it, resulting in a greater amount of plastids being inherited by the apical cell after cell division (Fig. 1b and Video S1).

**Figure 1.**
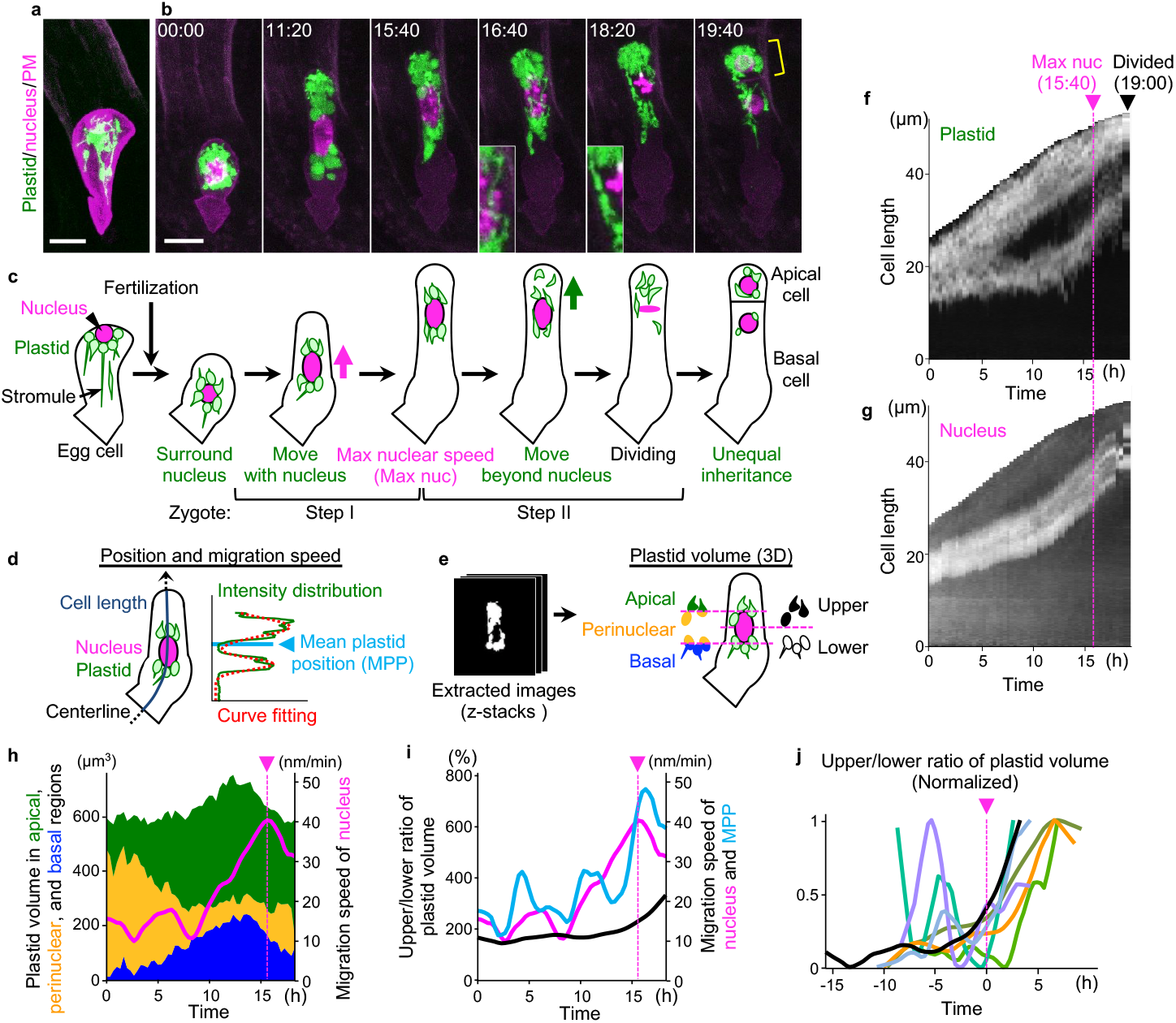
Plastids migrate apically in two steps and are unequally inherited by the apical cell. (**a** and **b**) Two-photon excitation microscopy (2PEM) images of the egg cell (**a**) and time-lapse images of the zygote in *in vitro*-cultivated ovules (**b**) expressing the plastid/nucleus/plasma membrane (PM) marker. Maximum intensity projection (MIP) images are shown, and numbers indicate the time (h:min) from the first frame. Yellow bracket indicates the apical cell, and the insets show magnified images of the perinuclear regions. (**c**) Illustrations summarising the respective stages. (**d** and **e**) Schematics of image quantification methods to detect the position and migration speed (**d**) and the volume (**e**) of plastids. For details, see Methods. (**f** and **g**) Kymographs showing the signal intensity of the plastid (**f**) and nucleus (**g**) projected onto the zygote centerline. Magenta and black arrowheads indicate the time points of maximum nuclear speed (Max nuc) and just after cell division (Divided), respectively. (**h** and **i**) Time courses of plastid volume in the apical (green), perinuclear (orange), and basal (blue) regions (**h**) and of the ratio of plastid volume of upper and lower regions (black) and the migration speed of mean plastid position (MPP; cyan) (**i**). Nuclear migration speed (magenta) is shown in both graphs. (**j**) Time course of the ratio of plastid volume of upper and lower regions (n = 7). Values were normalized with the minimum set to 0 and the maximum set to 1. Time is shown with each sample’s Max nuc point set to 0. The black line corresponds to the sample shown in (**i**). Scale bars: 10 µm.

To quantify these dynamics, plastid and nuclear signals were projected onto the cell centerline to determine the positions and velocities of the nucleus and the mean plastid position (MPP) (Fig. 1d), and plastid volumes were extracted from z-stacks to determine the amounts in the upper and lower regions, as well as in apical, perinuclear, and basal compartments relative to the nucleus (Fig. 1e) (Video S1, see Methods and Fig. S3 for details). During active nuclear migration (from 08:20 to 15:40), plastids moved while remaining in the upper and lower regions relative to the nucleus, showing high intensities above and below the nucleus in the kymographs (Fig. 1f and g) and maintaining low perinuclear volumes (Fig. 1h). After the time point of maximum nuclear speed (Max nuc; 15:40), that is, when the nucleus began to decelerate (Fig. 1h, magenta line), plastids passed the nucleus and advanced apically, as shown by an increase in perinuclear plastid volume (Fig. 1h), a higher plastid velocity than that of the nucleus (Fig. 1i, the cyan line exceeds the magenta line), and an increase in the upper/lower plastid-volume ratio (Fig. 1i, black line). This behavior was consistent across samples, with the upper/lower ratio increasing around the Max nuc time point (Fig. 1j). These observations show that plastids reach the apical region through two sequential steps, switching from Step I, in which plastids occupy upper and lower positions relative to the nucleus during active nuclear movement, to Step II, in which plastids pass the decelerating nucleus and move beyond the nucleus (Fig. 1c).

### The first step of plastid migration depends on F-actin and the nucleus, and the second step requires MTs

To identify the driving forces of plastid migration, we first assessed the involvement of F-actin, because we previously found that all of the nuclei, mitochondria, and vacuoles migrate along longitudinal F-actin arrays in zygotes ^3-5^. Treatment of zygotes expressing the plastid/nucleus/PM marker with latrunculin B (LatB), an inhibitor of actin polymerization, arrested nuclear migration as previously reported ^3^; however, plastids continued to migrate beyond the nucleus and accumulated apically (Fig. 2a and Video S2). Consequently, a greater amount of plastids was inherited by the apical cell, and in the extreme case (1 of 9 samples), all plastids were inherited by the apical cell (Fig. S1 and Video S3). These results indicate that F-actin is not essential for the unequal plastid inheritance after cell division, but rather is important for avoiding overly extreme segregation, such as all-or-none outcomes.

**Figure 2.**
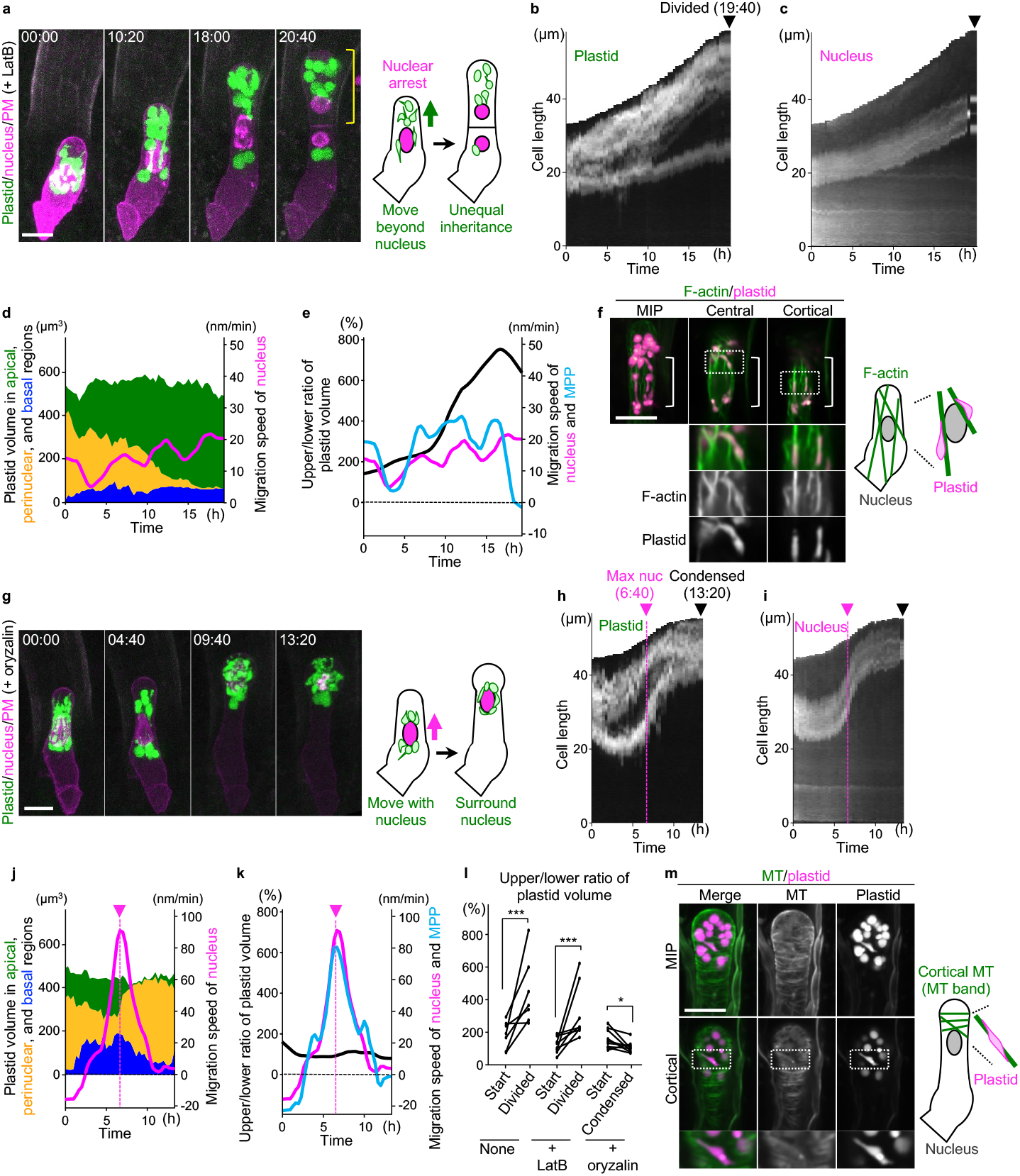
The two-step plastid migration depends on F-actin and MT, respectively. (**a**-**e, g**-**k**) Effects of treatment with the polymerization inhibitors for F-actin (1 μM LatB, **a**-**e**) and MTs (10 μM oryzalin, **g**-**k**). (**a** and **g**) Time-lapse 2PEM images of plastid/nucleus/PM marker. MIP images are shown, and numbers indicate the time (h:min) from the first frame. Yellow bracket indicates the apical cell. The illustrations on the right summarize the effects. (**b, c, h, i**) Kymographs showing the signal intensity of the plastid (**b** and **h**) and nucleus (**c** and **i**) projected onto the zygote centerline. Magenta arrowhead indicates the timing of Max nuc. Black arrowheads indicate the time point just after cell division (**b** and **c**) and nuclear condensation (**h** and **i**). (**d, e, j, k**) Time courses of plastid volume in the apical (green), perinuclear (orange), and basal (blue) region (**d** and **j**) and of the ratio of plastid volume of upper and lower regions (black) and the migration speed of MPP (cyan) (**e** and **k**). Nuclear migration speed (magenta) is shown in all graphs. (**f** and **m**) 2PEM images of F-actin/plastid (**f**) and MT/plastid (**m**) markers. MIP images (**f** and **m**) and single-plane images at the central (**f**) and cortical (**f** and **m**) regions are shown. Lower panels show magnified images of the boxed regions, and the bracket indicates the nuclear region. (**l**) The ratio of plastid volume of upper and lower regions in untreated (None), LatB-and oryzalin-treated zygotes. The difference between the observation start (Start) and just after cell division (Divided) is compared, except for the oryzalin-treated zygotes, for which the ratio at nuclear condensation (Condensed) was measured. Significance was determined by the Brunner–Munzel test (\**P* < 0.05 and \*\*\**P* < 0.001; n = 7, 8, and 9 for None, LatB- and oryzalin-treated zygotes, respectively). Note that the ratio decreased at Condensed compared with Start in oryzalin-treated zygotes. Scale bars: 10 µm.

Although Steps I and II could not be distinguished in LatB-treated zygotes because nuclear arrest prevented us from detecting the Max nuc time point (Fig. 2d, no typical peak in the magenta line), filamentous plastids passing along the lateral side of the nucleus to move beyond it were observed in the early elongation phase (6–12 h; Fig. 2a). During this period, perinuclear plastid volume was high (Fig. 2b–d), and plastid velocity exceeded that of the nucleus (Fig. 2e, the cyan line exceeds the magenta line), suggesting that plastids detached from the nucleus prematurely and entered Step II. Plastids colocalized with F-actin near the nucleus in a F-actin/plastid marker that contains a F-actin marker (EC1p::Lifeact-mNeonGreen) and a plastid marker (EC1p::TP-tdTomato) (Fig. 2f). Taken together with the above results from LatB-treated zygotes, plastids may be tethered to the nucleus via F-actin, as chloroplasts do during innate immune responses ^11^.

During innate immunity, chloroplasts also migrate along microtubules (MTs) ^11^. Treatment of zygotes with oryzalin, a MT-depolymerizing inhibitor that disrupts cortical MTs and the division apparatus, led to a swollen cell tip and failed nuclear division as previously reported (Fig. 2g and Video S4) ^3^. Despite these abnormalities, up to the Max nuc time point (06:40), plastids in oryzalin-treated zygotes moved properly with the nucleus while remaining above and below it (Fig. 2h–j, compare to Fig. 1f–h), indicating that MTs are not essential for Step I migration. After the Max nuc timing, however, plastids failed to detach from the nucleus and decelerated at the same rate as the nucleus (Fig. 2k, the cyan and magenta lines show a similar pattern). At the nuclear-condensation time point (13:20), which corresponds to the timing of cell division in untreated zygotes, plastids remained closely associated with the nucleus without any detectable asymmetry (Fig. 2g). Accordingly, the upper/lower plastid-volume ratio did not increase (Fig. 2k, black line), indicating that MTs are essential for Step II migration, during which plastids move beyond the nucleus and thereby acquire asymmetry. This conclusion was statistically supported: in untreated and LatB-treated zygotes, the upper/lower ratio increased significantly from the observation start (‘Start’) to just after cell division (‘Divided’), whereas in oryzalin-treated samples the ratio did not increase at the nuclear condensation timing (‘Condensed’), and instead slightly decreased (Fig. 2l).

In the MT/plastid marker that contains a MT marker (EC1p::Clover-TUA6) and the above-mentioned plastid marker (EC1p::TP-tdTomato), plastids colocalized with cortical MTs, which align transversely in the subapical region as the so-called subapical MT band (Fig. 2m) ^12^. Together, these results show that plastid migration switches from an F-actin–dependent Step I, in which plastids move with the nucleus, to an MT-dependent Step II, in which plastids move beyond the nucleus, thereby likely ensuring robust, biased plastid inheritance while avoiding both overly extreme outcomes and equal segregation.

### Starch granules within plastids are not required for polar migration

Plastid function varies according to its contents, as exemplified by chloroplasts that contain chlorophyll and amyloplasts that accumulate starch granules ^13^. We therefore examined the contents of plastids in the zygote. Consistent with previous observations ^10^, electron microscopy revealed spherical starch granules in the apical region of the egg cell (Fig. 3a). A similar distribution was observed using green-fluorescent probes that visualize starch in living plant cells, fluorescein-5-tert-butyl carbamate (Probe 3b) and fluorescein diacetate (FDA) ^14,15^, and several starch granules were detected within plastids labeled with the plastid marker (EC1p::TP-tdTomato) described above (Fig. S2a, b). Similar starch granules were also detected in zygotes (Fig. S2b), indicating that plastids in egg cells and zygotes are starch-containing amyloplasts.

**Figure 3.**
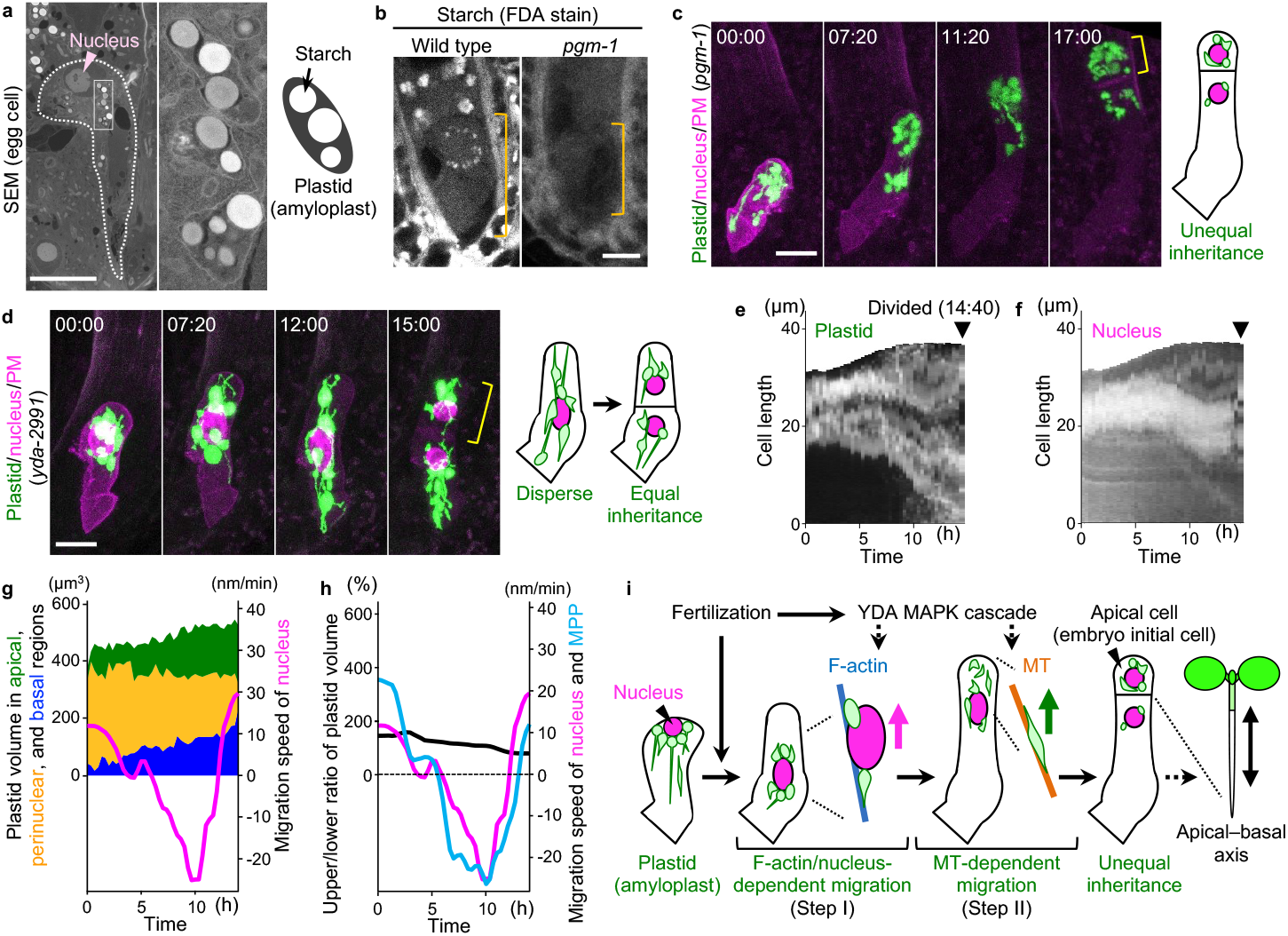
Polar plastid migration is not driven by starch granules, but depends on YDA pathway. (**a**) Scanning electron microscopy images of the egg cell. Magnified image of the boxed region is shown. The cell is outlined. (**b**) 2PEM images of wild type and *pgm-1* zygotes stained with a starch probe (fluorescein diacetate; FDA). Midplane images are shown, and the orange bracket indicates the zygotes. (**c** and **d**) Time-lapse 2PEM images of *pgm-1* and *yda-2991* zygotes expressing the plastid/nucleus/PM marker. MIP images are shown, and numbers indicate the time (h:min) from the first frame. Yellow bracket indicates the apical cell. The illustrations on the right summarize the effects. (**e** and **f**) Kymographs of *yda-2991* zygotes showing the signal intensity of the plastid (**e**) and nucleus (**f**) projected onto the zygote centerline. Black arrowheads indicate the time point just after cell division. (**g** and **h**) Time courses of plastid volume in the apical (green), perinuclear (orange), and basal (blue) region (**g**) and of the ratio of plastid volume of upper and lower regions (black) and the migration speed of MPP (cyan) (**h**). Nuclear migration speed (magenta) is shown in both graphs. (**i**) Schematic diagram summarizing the plastid behavior during apical–basal axis formation. Scale bars: 10 µm.

In roots and stems, amyloplasts sediment in the direction of gravity, thereby altering cell polarity and inducing gravitropism ^16^. We therefore examined the contribution of starch to polar plastid migration in the zygote using the *pgm-1* mutant, which is defective in the starch-biosynthetic enzyme PHOSPHOGLUCOMUTASE and consequently shows impaired gravitropism ^17^. In *pgm-1*, starch granules were detected in neither the zygote nor the surrounding endosperm and ovule tissues (Fig. 3b), as previously reported ^18^, and plastids in the egg cell were no longer spherical but appeared as thin fiber-like structures (Fig. S2c). Despite these starch-free plastids, *pgm-1* zygotes expressing the plastid/nucleus/PM marker showed no obvious defects in plastid migration with and beyond the nucleus, unequal plastid inheritance by the apical cell, zygote elongation, or asymmetric cell division (Fig. 3c and Video S5). Thus, although starch granule accumulation contributes to plastid morphology, it is not essential for polar plastid migration or zygote polarization. These findings suggest that, in the zygote, plastid behavior is controlled by a mechanism distinct from the simple physical response underlying gravitropism.

### Polar plastid migration depends on the YDA MAPKKK pathway

We therefore focused on a mitogen-activated protein kinase (MAPK) pathway that is activated in a fertilization-dependent manner and promotes zygote polarization ^19^. In order to examine its contribution to plastid behavior, we used the *yda-2991* mutant of YODA (MAPK kinase kinase), which fails to support directional movement of the nucleus and vacuoles along F-actin and exhibits premature termination of cell elongation accompanied by loss of the subapical MT band ^12,20^.

In *yda-2991* zygotes expressing the plastid/nucleus/PM marker, plastids initially surrounded the nucleus, and the zygote began to elongate as in the wild type (Fig. 3d and Video S6). Shortly thereafter, however, zygote elongation ceased, and the nucleus began to move back toward the cell center (Fig. 3e, f), as indicated by the negative nuclear velocity (Fig. 3g, magenta line). Although the Max nuc timing could not be defined (Fig. 3g, no positive peak in the magenta line), during the later period of zygote elongation, plastids did not move apically beyond the nucleus; instead, they became elongated while remaining associated with the nucleus and dispersed throughout the cell (Fig. 3d, e, g). Accordingly, the plastid and nucleus showed similar velocity (Fig. 3h, the cyan and magenta lines show a similar pattern), and the upper/lower plastid-volume ratio did not increase (Fig. 3h, black line), resulting in equal plastid inheritance. These results indicate a requirement for YDA in Step II migration, which drives plastids beyond the nucleus. Notably, plastids continuously associate with the nucleus, suggesting that YDA is not essential for plastid tethering to the nucleus during Step I, although YDA is required for the F-actin-dependent nuclear migration itself that underlies Step I migration of plastids.

## Discussion

In summary, we found that zygotic plastids undergo a two-step polar migration, switching from F-actin to MTs (Fig. 3i). This is different from other organelles previously examined in the Arabidopsis zygote, such as the nucleus, vacuoles, and mitochondria, which have been reported to rely primarily on F-actin for polar migration. The two-step polar migration is likely essential to ensure a properly balanced plastid distribution that supports subsequent embryogenesis, while avoiding both overly extreme outcomes and equal segregation. Plastids in zygotes contain starch granules, but these are not essential for plastid polar migration, unlike in gravitropism. Because the apical cell is an embryo initial cell that generates most of the plant body, preferential inheritance of starch, an important carbon source, by the apical cell at the time of zygotic division likely helps support the subsequent phase of active cell proliferation, although maternally supplied sugars also contribute to embryogenesis ^21^.

A two-step plastid migration has also been described in tobacco epidermal cells during immune responses, in which chloroplasts first move along MTs and are then anchored by perinuclear F-actin ^11^. In protonemal cells of *Physcomitrium patens*, chloroplasts can move along both F-actin and MTs ^22^. However, active dissociation of plastids from the nucleus followed by polar migration toward the cell periphery, with trafficking sequentially along F-actin and MTs, has not yet been reported, suggesting that zygotes may employ a distinct mechanism. In particular, plastids in the zygote would require a mechanism to switch from Step I to Step II, whereby they dissociate from the nucleus when its movement decelerates, and then transfer to MTs to reach the apical tip. Recent studies have proposed that zygote elongation increases surface tension in the subapical region, which guides cortical MT polymerization and results in the continuous maintenance of the subapical cortical MT band during zygote growth ^12,23^. Because this MT band is continuously repositioned apical to the nucleus, it is reasonable that plastids moving beyond the nucleus utilize MTs. To elucidate the molecular mechanism underlying this two-step migration, future identification of downstream components of the YDA pathway will be helpful. For example, this effort may reveal regulators analogous to KINESIN REQUIRED FOR INDUCING STROMULES 1 (KIS1), which contains an MT-binding motor domain and an actin-binding domain and regulates the chloroplast movement and anchoring during immune responses ^24^, and modifiers that regulate their activity in coordination with cell-cycle progression. Such discoveries should help provide a more comprehensive understanding of how fertilization-triggered signals are translated into the unequal inheritance of plastids.

## Methods

### Plant material and growth conditions

All Arabidopsis lines were in the Columbia (Col-0) background. *pgm-1* and *yda-2991* have been described previously ^20,25^. The nuclear marker in the plastid/nucleus/PM marker is ABI4p::H2B-tdTomato (coded as MU2463), which expresses HISTONE H2B fused to a red fluorescent protein tdTomato under the control of the *ABA INSENSITIVE4* promoter, as described previously ^12^. The F-actin marker (EC1p::Lifeact-mNeonGreen; coded as MU2232) contains a green-fluorescent protein mNeonGreen fused to the Lifeact sequence under the control of the *EGG CELL1* (*EC1*) promoter ^5^. The MT marker (EC1p::Clover-TUA6; coded as MU2228) contains a green-fluorescent protein Clover fused with TUBULIN ALPHA6 under the control of the *EC1* promoter ^26^.

Plants were grown at 20–24°C under continuous light or long-day conditions (16-h-light/8-h-dark photoperiod).

### Plasmid construction

The plastid/nucleus/PM marker contains EC1p::TP-Clover as a plastid marker, the above-mentioned ABI4p::H2B-tdTomato as nuclear marker, and EC1p::RRvT-SYP121 as a PM marker. In EC1p::TP-Clover (coded as MU2381), the 463-bp *EC1* promoter ^27^ was fused to a part of 3’UTR (75 bp) and the coding region corresponding to the transit peptide to plastid (TP, 1-90 amino acids) of FTSZ1 (AT5G55280), Clover protein, and NOPALINE SYNTHASE (NOS) terminator in a pMDC99 binary vector ^28^. In EC1p::RRvT-SYP121 (coded as YK80), the 463-bp *EC1* promoter was fused to the coding region of red fluorescent protein RRvT ^29^, SYNTAXIN RELATED PROTEIN121 (AT3G11820), and the NOS terminator in the pPZP211 binary vector ^30^. The plastid marker showing red fluorescence contains EC1p::TP-tdTomato (coded as MU2328), in which the 463-bp *EC1* promoter was fused to the same part of FTSZ1 used in EC1p::TP-Clover, tdTomato, and the NOS terminator in pSMAB704 binary vector ^31^.

These constructs were transformed into Arabidopsis using the floral dip method ^32^.

### Time-lapse observation and histological analysis

The *in vitro* ovule cultivation for zygote live-cell imaging was performed using the Nitsch medium, as previously described ^33,34^. 2PEM images were all acquired using a laser-scanning inverted microscope (A1; Nikon) equipped with a pulse laser (Mai Tai DeepSee; Spectra-Physics) or a laser-scanning inverted microscope (AX; Nikon) equipped with an ultrafast tunable laser (InSight X3 Dual option; Spectra-Physics). Fluorescence signals were detected by the external non-descanned GaAsP PMT detectors. For the A1 system, two dichroic mirrors (DM495 and DM560) were used, along with a band-pass filter (525/50 nm) for green signals (Clover, NeonGreen, FDA, and Probe 3b), and a mirror for red signals (tdTomato and RRvT). For the AX system, three dichroic mirrors (DM488, DM560 and DM685) were used, along with two band-pass filters (525/50 nm for green, and 605/70 nm for red). Time-lapse images were acquired every 20 min as 35 *z*-stacks with 1.0-μm intervals using a 40× water-immersion objective lens (CFI Apo LWD WI; Nikon) with Immersol W 2010 (Zeiss) immersion medium.

For inhibitor treatment, individual inhibitors [0.1% DMSO containing 10 μM oryzalin (Chem service #N-12729) or 1 μM LatB (Funakoshi #10-4303)] were added to the cultivation medium approximately 1 h before observation, as previously described ^35,36^.

For starch staining, 10 μM FDA (Tokyo Chemical Industry # F0240) or 5 μM Probe 3b ^15^ was added to the cultivation medium approximately 1 h before observation.

For the analysis of plastid association with F-actin and MT, the snapshot images were processed with “Denoise” function in NIS-Elements software (Nikon).

Maximum intensity projection (MIP) and image processing were performed using NIS-Elements software, or Fiji (https://fiji.sc/).

### Quantification of nuclear and plastid dynamics from live-cell imaging sequence

The analysis workflow is summarized in Fig. S3. From raw time-lapse z-stacks (Fig. S3a), we generated two data streams: a MIP image for 2D analyses (Fig. S3b–j) and the green-channel z-stacks for 3D plastid volumetry (Fig. S3k–p).

For the 2D analyses, we extracted the magenta channel from the MIP (Fig. S3c) and obtained binary masks for the PM and nucleus (Fig. S3d and g, respectively). From the PM mask, we extracted the cell contour (Fig. S3e) and the centerline (Fig. S3f) by following the KymoTip pipeline, as described previously ^37^. We resampled the centerline into equally spaced points and computed the normal vector at each point. For each normal vector, we subdivided the in-cell segment into 100 sampling points, sampled fluorescence intensities, and mapped the mean intensity back onto the corresponding centerline position. Repeating this procedure over time frames yielded kymographs for the nuclear and plastid signals.

For nuclear dynamics, we constructed a one-dimensional nuclear intensity profile along the centerline and normalized it to 0–1. We then fitted a Gaussian function and defined the fitted peak position as the nuclear center measured from the basal end of the centerline (Fig. S3h). From time-lapse data, nuclear migration speed was computed using a central-difference scheme and smoothed using LOWESS ^38^.

For plastid position dynamics, we extracted the green channel from the MIP (Fig. S3i), generated a one-dimensional plastid intensity profile along the centerline, and fitted it with the sum of two Gaussian functions. We defined the MPP as the amplitude-weighted average of the two fitted peak positions (Fig. S3j). Plastid migration speed was computed from the MPP time series using central differences and smoothing as above.

For plastid volumetry, we obtained the green-channel z-stacks (Fig. S3k) and generated plastid binary images by Otsu thresholding (Fig. S3l). For two-region measurements, we determined the nuclear centroid and split the plastid image into upper and lower halves at the nuclear center (Fig. S3m), measured plastid volumes in each half, and calculated the upper/lower plastid volume ratio (Fig. S3n). For three-region measurements, we used the nuclear bounding box to define nuclear upper and lower edges, partitioned the cell into apical, perinuclear, and basal regions (Fig. S3o), and quantified plastid volumes for each region (Fig. S3p). Volume ratios were smoothed as above.

### Electron microscope observation

Semi-thin sections were observed using a scanning electron microscope as follows. Wild-type ovules were fixed in a fixation solution (mixture of 2% glutaraldehyde and 4% paraformaldehyde in 50 mM sodium cacodylate buffer, pH 7.4), overnight at 4°C. The tissue segments were washed in the buffer and post-fixed in 2% osmium tetroxide in the buffer for 3 h at 4°C. The tissue was then dehydrated through a graded ethanol series, incubated in ethanol, transferred to propylene oxide, infiltrated, and embedded in resin (Quetol 651, Nissin-EM). The resin blocks were cut into semi-thin sections (500 nm thick) using a diamond knife with a large water trough (Histo-Jumbo, DiATOME) mounted on an ultramicrotome (EM UC6, Leica), as described previously ^39^. Serial sections collected on glass slides were stained with an alternative uranium acetate aqueous (EM Stainer, Nissin-EM) for 15 min, followed by 0.4% lead citrate (TAAB) aqueous for 4 min, and coated with gold for 7 sec using an ion-sputtering apparatus (IB-3, Eiko). The sections were observed with a field emission scanning electron microscope (SU8600, Hitachi), at an accelerating voltage of 5 kV.

### Quantification and statistical analysis

Statistical analysis was performed using R. The methods used for the statistics and the number of samples are described in the respective legend.

## Acknowledgments

We thank Hisa Yoshida, Tamiko Ambo, Satomi Watanabe, Yuko Kudo and Junko Kato for technical support, Hiroaki Ichikawa and Makoto Fujiwara for providing pSMAB704 binary vector, and Miyo Terao Morita for *pgm-1*.

This work was supported by the Japan Society for the Promotion of Science [a Grant-in-Aid for Early-Career Scientists (JP22K15135 and JP25K18484 to H.M., JP23K14204 to Y.Kimata, and JP25K18499 to Z.K.), Grant-in-Aid for Transformative Research Areas (A) (JP25H01809 to Y.Kimata; JP20H05905 and JP25H01340 to Y.Kodama), a Grant-in-Aid for Scientific Research on Innovative Areas (JP16H06280 (Advanced Bioimaging Support)), a Grant-in-Aid for Scientific Research (B) (JP23H01143 to S.T. and JP23H02494 to M.U.), and International Leading Research (JP22K21352) to M.U.], the Japan Science and Technology Agency [CREST (JPMJCR2121 to S.T. and M.U., YORC to H.M., Y.Kimata, Z.K. and T.N.)], the Inamori Foundation (Inamori Research Grant to Y.Kimata), and the Suntory Rising Stars Encouragement Program in Life Sciences (SunRiSE to M.U.).

## Author Contributions

M.U. designed the research; K.T., H.M., T.O., S.K., S.H., S.I., and Y.Kodama carried out the experiments; Z.K., T.N., S.T., and Y.Kimata analyzed data; and K.T. and M.U. wrote the manuscript.

## Data availability

Data supporting the findings of this work are available within the paper and its Supplementary files. Plasmids and transgenic lines generated in this study are available upon reasonable request to the corresponding author.

## Code availability

No new code was generated in this paper. All analytical methods are described in the Methods section.

## Supplementary materials

Figures S1 to S3

Videos S1 to S6

## References

1. Mansfield, S.G. & Briarty, L.G. Early embryogenesis in Arabidopsis thaliana. II. The developing embryo. Can J Bot 69, 461–476 (1991).

2. Tanaka, S. et al. HD-ZIP IV genes are essential for embryo initial cell polarization and the radial axis formation in Arabidopsis. Current Biology 34, 4639-4649.e4 (2024).

3. Kimata, Y. et al. Cytoskeleton dynamics control the first asymmetric cell division in Arabidopsis zygote. Proc Natl Acad Sci U S A 113, 14157–14162 (2016).

4. Kimata, Y. et al. Mitochondrial dynamics and segregation during the asymmetric division of Arabidopsis zygotes. Quantitative Plant Biology 1, e3 (2020).

5. Kimata, Y. et al. Polar vacuolar distribution is essential for accurate asymmetric division of Arabidopsis zygotes. Proc Natl Acad Sci U S A 116, 2338–2343 (2019).

6. Jensen, W.A. Cotton embryogenesis: The zygote. Planta 79, 346–66 (1968).

7. Suzuki, K., Taniguchi, T. & Maeda, E. Ultrastructure and Cleavage Pattern of Rice Proembryos. Japanese Journal of Crop Science 61, 292–303 (1992).

8. Baldwin, A. et al. A molecular-genetic study of the Arabidopsis Toc75 gene family. Plant Physiol 138, 715–33 (2005).

9. Kovacheva, S., Bedard, J., Wardle, A., Patel, R. & Jarvis, P. Further in vivo studies on the role of the molecular chaperone, Hsp93, in plastid protein import. Plant J 50, 364–79 (2007).

10. Mansfield, S.G., Briarty, L.G. & Erni, S. Early embryogenesis in Arabidopsis thaliana. I. The mature embryo sac. Can J Bot 69, 447–460 (1991).

11. Kumar, A.S. et al. Stromule extension along microtubules coordinated with actin-mediated anchoring guides perinuclear chloroplast movement during innate immunity. Elife 7(2018).

12. Kang, Z. et al. Temporal changes in surface tension guide the accurate asymmetric division of Arabidopsis zygotes. bioRxiv, doi: 10.1101/2024.08.07.605794 (preprint) (2024).

13. Choi, H., Yi, T. & Ha, S.H. Diversity of Plastid Types and Their Interconversions. Front Plant Sci 12, 692024 (2021).

14. Ichikawa, S., Sakata, M., Oba, T. & Kodama, Y. Fluorescein staining of chloroplast starch granules in living plants. Plant Physiol 194, 662–672 (2024).

15. Kusano, S., Nakamura, S., Izumi, M. & Hagihara, S. A fluorescent molecular probe for live-cell imaging of starch granules. ChemRxiv (2023).

16. Morita, M.T. & Tasaka, M. Gravity sensing and signaling. Current Opinion in Plant Biology 7, 712–718 (2004).

17. Caspar, T. & Pickard, B.G. Gravitropism in a starchless mutant of Arabidopsis : Implications for the starch-statolith theory of gravity sensing. Planta 177, 185–97 (1989).

18. Hedhly, A. et al. Starch Turnover and Metabolism during Flower and Early Embryo Development. Plant Physiol 172, 2388–2402 (2016).

19. Ueda, M. et al. Transcriptional integration of paternal and maternal factors in the Arabidopsis zygote. Genes Dev 31, 617–627 (2017).

20. Matsumoto, H., Kimata, Y., Higaki, T., Higashiyama, T. & Ueda, M. Dynamic Rearrangement and Directional Migration of Tubular Vacuoles are Required for the Asymmetric Division of the Arabidopsis Zygote. Plant Cell Physiol 62, 1280–1289 (2021).

21. Chen, L.Q. et al. A cascade of sequentially expressed sucrose transporters in the seed coat and endosperm provides nutrition for the Arabidopsis embryo. Plant Cell 27, 607–19 (2015).

22. Sato, Y., Wada, M. & Kadota, A. Choice of tracks, microtubules and/or actin filaments for chloroplast photo-movement is differentially controlled by phytochrome and a blue light receptor. Journal of Cell Science 114, 269–279 (2001).

23. Nonoyama, T. et al. Agent-based simulation of cortical microtubule band movement in arabidopsis zygotes. Sci Rep 15, 25787 (2025).

24. Meier, N.D., Seward, K., Caplan, J.L. & Dinesh-Kumar, S.P. Calponin homology domain containing kinesin, KIS1, regulates chloroplast stromule formation and immunity. Science Advances 9, eadi7407 (2023).

25. Caspar, T., Huber, S.C. & Somerville, C. Alterations in Growth, Photosynthesis, and Respiration in a Starchless Mutant of Arabidopsis thaliana (L.) Deficient in Chloroplast Phosphoglucomutase Activity. Plant Physiol 79, 11–7 (1985).

26. Horiuchi, R. et al. Deep learning-based cytoskeleton segmentation for accurate high-throughput measurement of cytoskeleton density. Protoplasma (2024).

27. Sprunck, S. et al. Egg cell-secreted EC1 triggers sperm cell activation during double fertilization. Science 338, 1093–7 (2012).

28. Curtis, M.D. & Grossniklaus, U. A gateway cloning vector set for high-throughput functional analysis of genes in planta. Plant Physiol 133, 462–9 (2003).

29. Wiens, M.D. et al. A Tandem Green–Red Heterodimeric Fluorescent Protein with High FRET Efficiency. ChemBioChem 17, 2361–2367 (2016).

30. Hajdukiewicz, P., Svab, Z. & Maliga, P. The small, versatile pPZP family of Agrobacterium binary vectors for plant transformation. Plant Mol Biol 25, 989–94 (1994).

31. Igasaki, T., Ishida, Y., Mohri, T., Ichikawa, H. & Shinohara, K. Transformation of Populus alba and direct selection of transformants with the herbicide Bialaphos. Bulletin of FFPRI 1, 235–240 (2002).

32. Clough, S.J. & Bent, A.F. Floral dip: a simplified method for Agrobacterium-mediated transformation of Arabidopsis thaliana. Plant J 16, 735–43 (1998).

33. Kurihara, D., Kimata, Y., Higashiyama, T. & Ueda, M. In Vitro Ovule Cultivation for Live-cell Imaging of Zygote Polarization and Embryo Patterning in Arabidopsis thaliana. J Vis Exp (2017).

34. Ueda, M., Kimata, Y. & Kurihara, D. Live-Cell Imaging of Zygotic Intracellular Structures and Early Embryo Pattern Formation in Arabidopsis thaliana. Methods Mol Biol 2122, 37–47 (2020).

35. Hanaki, Y. et al. A simple and versatile plasma membrane staining method for visualizing living cell morphology in reproductive tissues across diverse plant species. Plant Methods 21, 149 (2025).

36. Kimata, Y. et al. Novel inhibitors of microtubule organization and phragmoplast formation in diverse plant species. Life Sci Alliance 6(2023).

37. Kang, Z. et al. KymoTip: High-throughput Characterization of Tip-growth Dynamics in Plant Cells. bioRxiv, 2025.06.27.661917 (2025).

38. Cleveland, W.S. Robust Locally Weighted Regression and Smoothing Scatterplots. Journal of the American Statistical Association 74, 829–836 (1979).

39. Ouk, R., Oi, T. & Taniguchi, M. Three-dimensional anatomy of mesophyll cells in rice leaf tissue by serial section light microscopy. Plant Production Science 23, 149–159 (2020).

