## Supplementary materials for "Two-step polar plastid migration via F-actin and microtubules ensures unequal inheritance during asymmetric division of Arabidopsis zygote"

**Supplementary materials include:**
Figures S1 to S3
Videos S1 to S6

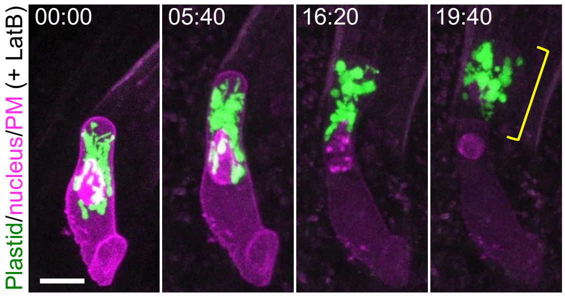


**Fig. S1. Effects of treatment with the F-actin polymerization inhibitor.**
Time-lapse 2PEM images of plastid/nucleus/PM marker in the presence of 1 μM LatB. MIP images are shown, and numbers indicate the time (h:min) from the first frame. Yellow bracket indicates the apical cell. Note that all plastids were inherited by the apical cell. Scale bar: 10 μm.

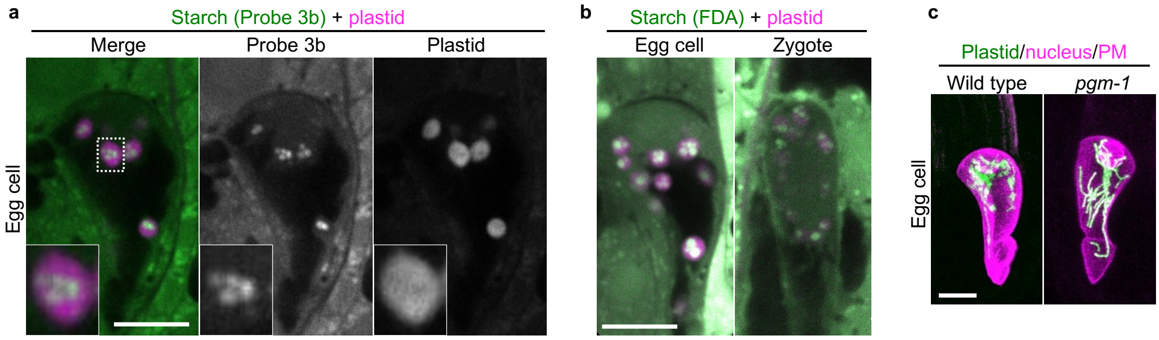


**Fig. S2. Plastids in the egg cells and zygotes are amyloplasts containing starch granules.** (**a** and **b**) 2PEM images of the egg cells (**a** and **b**) and the zygote (**b**) expressing a red-fluorescent plastid marker (EC1p::TP-tdTomato), stained with green-fluorescent starch probes (Probe 3b (**a**) and FDA (**b**)). Midplane images are shown, and magnified image of the boxed region is shown. (**c**) 2PEM images of the egg cells of wild type and *pgm-1* expressing the plastid/nucleus/PM marker. MIP images are shown. Scale bars: 10 μm.

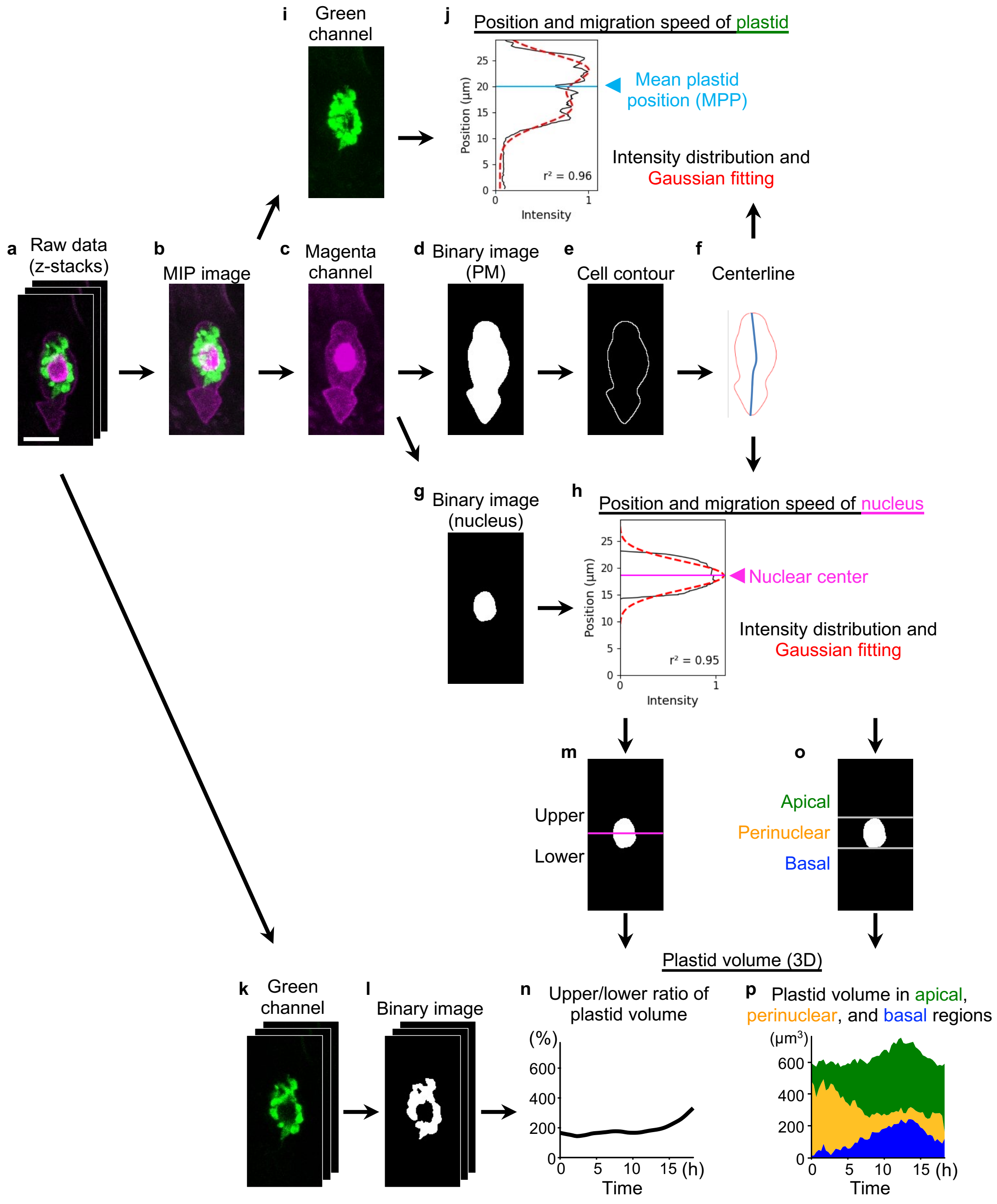


**Fig. S3. Procedure of image processing and quantification analysis.**
(**a**) Raw time-lapse z-stacks. (**b**) Maximum-intensity projection (MIP) image generated from the z-stacks. (**c**) Magenta-channel image extracted from the MIP, containing plasma membrane (PM) and nuclear signals. (**d**) Binary PM mask. (**e**) Cell contour extracted from the PM mask. (**f**) Centerline derived from the contour for one-dimensional measurements. (**g**) Binary nuclear mask. (**h**) Estimation of nuclear position from the centerline-based nuclear intensity distribution after Gaussian fitting; the fitted peak defines nuclear center (magenta arrowhead). (**i**) Green-channel image extracted from the MIP. (**j**) Estimation of plastid position from the centerline-based plastid intensity distribution after fitting with the sum of two Gaussian functions; the mean plastid position (MPP; cyan arrowhead) is computed as the amplitude-weighted average of the fitted peak positions. (**k**) Green-channel z-stack used for 3D plastid volumetry. (**l**) Binary plastid image after thresholding. (**m**) Schematic partitioning into upper and lower halves based on the nuclear center

for two-region plastid volumetry. (**n**) Example time course of the upper/lower plastid volume ratio. (**o**) Schematic partitioning into apical, perinuclear, and basal regions based on nuclear boundaries for three-region plastid volumetry. (**p**) Example time courses of plastid volume quantified in apical, perinuclear, and basal regions. Scale bar: 10 μm.

**Legend for Supplementary Videos**

**Video S1. Dynamics of plastid distribution in wild-type zygote.**
(Left and middle) Two-photon excitation microscopy (2PEM) time-lapse images of a wild-type zygote expressing the plastid (green), nuclear (magenta), and PM (magenta) marker. Numbers indicate the time (h:min) from the first frame. MIP images of merged channels (top left), green channel (top middle), and magenta channel (bottom left), and the extracted nuclear images (bottom middle) are shown. Mean plastid position (MPP) and nuclear center are shown as cyan and magenta dots, respectively. (Right) Signal distributions of plastids (top) and nucleus (bottom) projected onto the zygote centerline are shown. The dashed red lines indicate the signal distribution profile extracted by Gaussian fitting. MPP and nuclear center are shown as cyan and magenta bars, respectively. Coefficient of determination ($r^2$) at each time point is shown. Scale bar: 10 μm.

**Video S2. Dynamics of plastid distribution in the Latrunculin B-treated zygote.**
(Left and middle) 2PEM time-lapse images of a LatB-treated zygote expressing the plastid (green), nuclear (magenta), and PM (magenta) marker. Numbers indicate the time (h:min) from the first frame. MIP images of merged channels (top left), green channel (top middle), and magenta channel (bottom left), and the extracted nuclear images (bottom middle) are shown. MPP and nuclear center are shown as cyan and magenta dots, respectively. (Right) Signal distributions of plastids (top) and nucleus (bottom) projected onto the zygote centerline are shown. The dashed red lines indicate the signal distribution profile extracted by Gaussian fitting. MPP and nuclear center are shown as cyan and magenta bars, respectively. Coefficient of determination ($r^2$) at each time point is shown. Scale bar: 10 μm.

**Video S3. Excessively asymmetric plastid distribution in the Latrunculin B-treated zygote.**
2PEM time-lapse images of a LatB-treated zygote expressing the plastid (green), nuclear (magenta), and PM (magenta) marker. Numbers indicate the time (h:min) from the first frame. MIP images are shown. Scale bar: 10 μm.

**Video S4. Arrested plastid migration in the oryzalin-treated zygote.**
(Left and middle) 2PEM time-lapse images of an oryzalin-treated zygote expressing the plastid (green), nuclear (magenta), and PM (magenta) marker. Numbers indicate the time (h:min) from the first frame. MIP images of merged channels (top left), green channel (top middle), and magenta channel (bottom left), and the extracted nuclear images (bottom middle) are shown. MPP and nuclear center are shown as cyan and magenta dots, respectively. (Right) Signal distributions of plastids (top) and nucleus (bottom) projected onto the zygote centerline are shown. The dashed red lines indicate the signal distribution profile extracted by Gaussian fitting. MPP and nuclear center are shown as cyan and magenta bars, respectively. Coefficient of determination ($r^2$) at each time point is shown. Scale bar: 10 μm.

**Video S5. Proper plastid distribution in the *pgm-1* zygote.**
2PEM time-lapse images of a *pgm-1* zygote expressing the plastid (green), nuclear (magenta), and PM (magenta) marker. Numbers indicate the time (h:min) from the first frame. MIP images are shown. Scale bar: 10 μm.

**Video S6. Failure of plastid distribution in the *yda-2991* zygote.**
(Left and middle) 2PEM time-lapse images of a *yda-2991* zygote expressing the plastid (green), nuclear (magenta), and PM (magenta) marker. Numbers indicate the time (h:min) from the first frame. MIP images of merged channels (top left), green channel (top middle), and magenta channel (bottom left), and the extracted nuclear images (bottom middle) are shown. MPP and nuclear center are shown as cyan and magenta dots, respectively. (Right) Signal distributions of plastids (top) and nucleus (bottom) projected onto the zygote centerline are shown. The dashed red lines indicate the signal distribution profile extracted by Gaussian fitting. MPP and nuclear center are shown as cyan and magenta bars, respectively. Coefficient of determination ($r^2$) at each time point is shown. Scale bar: 10 μm.
